# Water-soluble tocopherol derivatives inhibit SARS-CoV-2 RNA-dependent RNA polymerase

**DOI:** 10.1101/2021.07.13.449251

**Authors:** Hayden T. Pacl, Jennifer L. Tipper, Ritesh R. Sevalkar, Andrew Crouse, Camerron Crowder, UAB Precision Medicine Institute, Shama Ahmad, Aftab Ahmad, Gillian D. Holder, Charles J. Kuhlman, Krishna C. Chinta, Sajid Nadeem, Todd J. Green, Chad M. Petit, Adrie J.C. Steyn, Matthew Might, Kevin S. Harrod

## Abstract

The recent emergence of a novel coronavirus, SARS-CoV-2, has led to the global pandemic of the severe disease COVID-19 in humans. While efforts to quickly identify effective antiviral therapies have focused largely on repurposing existing drugs^1–4^, the current standard of care, remdesivir, remains the only authorized antiviral intervention of COVID-19 and provides only modest clinical benefits^5^. Here we show that water-soluble derivatives of α-tocopherol have potent antiviral activity and synergize with remdesivir as inhibitors of the SARS-CoV-2 RNA-dependent RNA polymerase (RdRp). Through an artificial-intelligence-driven in silico screen and in vitro viral inhibition assay, we identified D-α-tocopherol polyethylene glycol succinate (TPGS) as an effective antiviral against SARS-CoV-2 and β-coronaviruses more broadly that also displays strong synergy with remdesivir. We subsequently determined that TPGS and other water-soluble derivatives of α-tocopherol inhibit the transcriptional activity of purified SARS-CoV-2 RdRp and identified affinity binding sites for these compounds within a conserved, hydrophobic interface between SARS-CoV-2 nonstructural protein 7 and nonstructural protein 8 that is functionally implicated in the assembly of the SARS-CoV-2 RdRp^6^. In summary, we conclude that solubilizing modifications to α-tocopherol allow it to interact with the SARS-CoV-2 RdRp, making it an effective antiviral molecule alone and even more so in combination with remdesivir. These findings are significant given that many tocopherol derivatives, including TPGS, are considered safe for humans, orally bioavailable, and dramatically enhance the activity of the only approved antiviral for SARS-CoV-2 infection^7–9^.

## Introduction

The emergence of multiple coronaviruses (CoVs) into the human population in recent years^10–12^ highlights the need for the rapid development of broad-spectrum interventional strategies against genetically distinct CoVs. Despite the unprecedented rapid development of effective vaccines targeting the severe acute respiratory syndrome coronavirus 2 (SARS-CoV-2), the etiologic agent of COVID-19, breakthrough infections in the immunocompromised^13–19^, infections in the young and elderly lacking a full repertoire of immunological responses^20^, vaccine reticence or hesitancy^21^, and the inequitable distribution of vaccines globally^22^ indicate that therapeutic interventions will be required to limit severe disease and substantial mortality associated with pandemic SARS-CoV-2 infection. Currently, remdesivir, under emergency FDA authorization, is the only antiviral to show improvement of clinical outcomes, albeit with limited effectiveness and the limitation of intravenous administration^5^. Furthermore, breakthrough infections and viral evolution of resistant strains is possible^14, 23^. This highlights the need for identifying new antivirals with broad efficacy against CoVs, and the potential for combinatorial interventional antivirals that reduce the development of resistance. Here, we employed a novel artificial intelligence query of FDA-approved compounds for repurposing as antivirals against SARS-CoV-2 and identify tocopherol derivatives with potent antiviral activity and synergy with remdesivir.

### Drug-repurposing against SARS-CoV-2

To select candidates for in vitro screening, we employed the open-source artificial intelligence software mediKanren to perform a logical query of the existing literature for compounds that are known to induce or inhibit host proteins involved in reducing or enhancing viral replication, respectively (https://github.com/webyrd/mediKanren) (Fig S1).

Complemented by a manual review of the available literature, we identified over 1300 compounds with proposed antiviral activity, of which nearly 100 were prioritized for in vitro screening (Table S1). To assess the antiviral activity of these compounds against virulent SARS-CoV-2, we adapted a traditional focus-forming assay in BSL-3 biocontainment by allowing foci to expand gradually over 48 hours, using progression of viral staining as a proxy for the magnitude of infection (Figure S2). Employing this method, we identified 12 compounds capable of reducing SARS-CoV-2 propagation by more than 90% at a concentration of 10 µM (Figure 1A). Follow-up testing revealed these compounds exhibit reliable dose-response relationships within our antiviral assay that are independent of their toxicity profile (Figure 1B).

**Figure 1.**
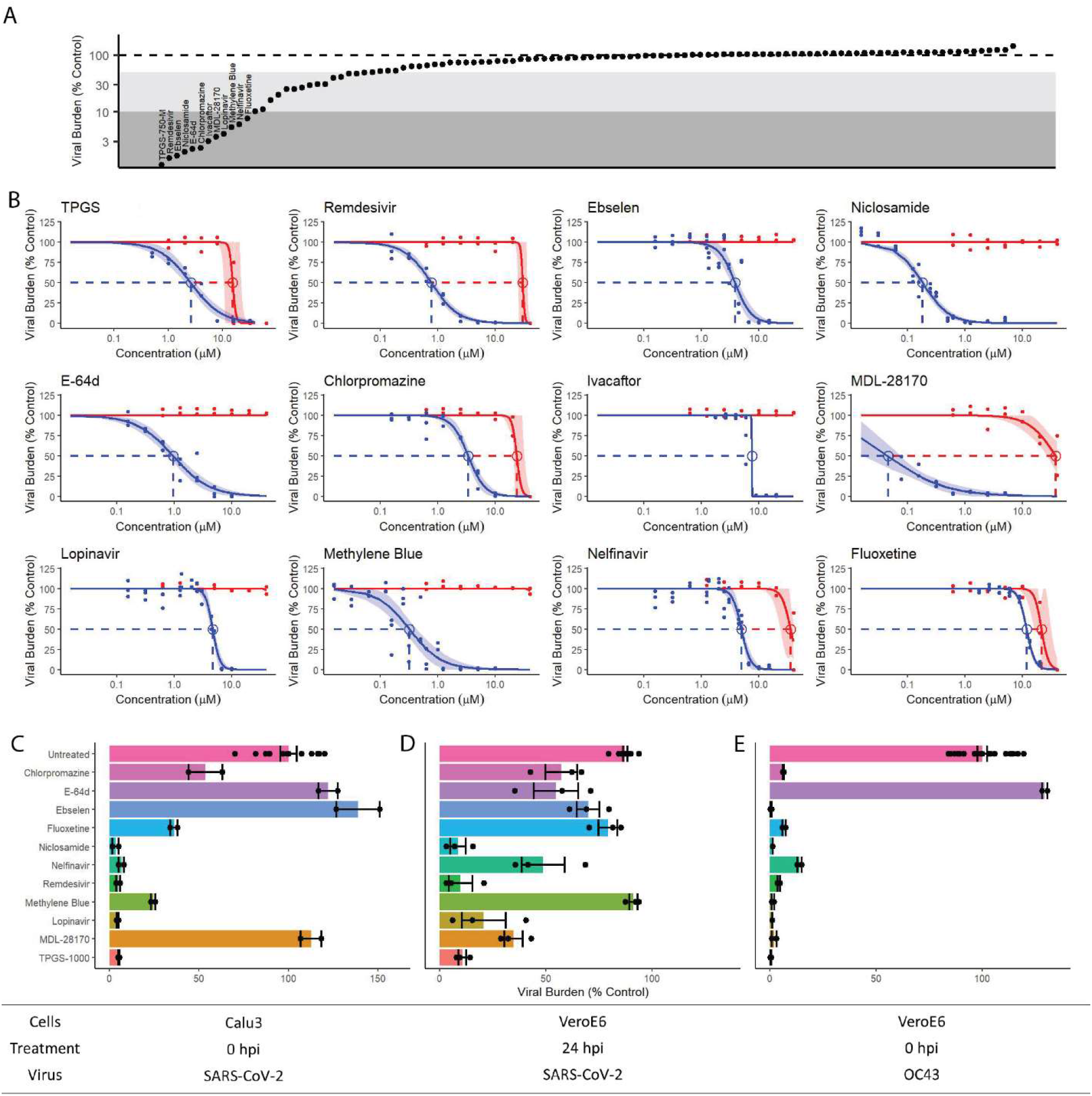
Identification and characterization of twelve FDA-approved compounds with antiviral activity against SARS-CoV-2. A) A graph depicting rank-ordered antiviral activity against SARS-CoV-2 at 10 µM for all drugs tested. Points represent the mean value from two independent experiments. B) Dose response curves for antiviral activity (blue) and viability (red) in VeroE6 cells. The shaded region around each curve represents the 95% confidence interval, while dots represent technical replicates pooled from two independent experiments. C-E) Antiviral activity of the top drugs at 10 µM in C) SARS-CoV-2-infected Calu3 cells, D) SARS-CoV-2-infected VeroE6 cells treated from 24 hpi until 48 hpi, and E) OC43-infected VeroE6 cells. Dots represent technical replicates from one experiment, bars and error bars represent the mean plus or minus the standard error of the mean. All experiments performed at least twice.

We expanded the characterization of candidate antiviral compounds by evaluation of antiviral activity across a range of conditions. We found that five of our top 12 compounds reduced the burden of SARS-CoV-2 in infected Calu3 cells, a human lung epithelial cell line, by greater than 90% (Figure 1C). Interestingly, four of these five drugs—niclosamide, remdesivir, lopinavir, and D-α-tocopherol polyethylene glycol succinate (TPGS)—were the only drugs to show maximum efficacy in halting an established SARS-CoV-2 infection in VeroE6 cells (Figure 1D). On the other hand, each of the top compounds except E-64d demonstrated strong antiviral activity against the seasonal β-coronavirus, OC43 (Figure 1E; Figure S3), suggesting that the majority of the compounds we identified are effective against β-coronaviruses more broadly. Overall, these findings describe four compounds with robust, broad-spectrum antiviral activity and therapeutic applications.

### TPGS synergizes with remdesivir

Given the threat of drug-resistance evolving in response to single-agent therapy, as well as the modest effects and limited supply of currently available antivirals for the treatment of COVID-19, we screened our top compounds for synergistic combinations. Relying on the principle of Loewe Additivity ^24, 25^, we combined our top compounds in a pairwise fashion to yield equipotent mixtures with a predicted efficacy of 50%. While the myriad combinations revealed mostly additive effects, the combination of remdesivir and TPGS resulted in an 8-fold increase in potency (Figure 2A). Follow-up testing of equipotent combinations of the top drugs with either remdesivir (Figure 2B; Figure S4) or TPGS (Figure 2C) revealed that this combination was indeed unique and dose-responsive across a range of concentrations. To fully characterize the synergy between TPGS and remdesivir we expanded our analysis across a range of mixtures to perform response-surface analysis, an extension of traditional isobologram analysis^25^. We identified that an equipotent mixture of TPGS and remdesivir was optimal for antiviral activity (Figure 2D, E), and, at this ratio, we observed a reduction of approximately 10-fold in the IC_90_ (Figure 2E). Of note, including one drug as at least 10% of the total combined dose dramatically enhances the potency of the other, suggesting that synergism can be honed to limiting availability to either compound.

**Figure 2.**
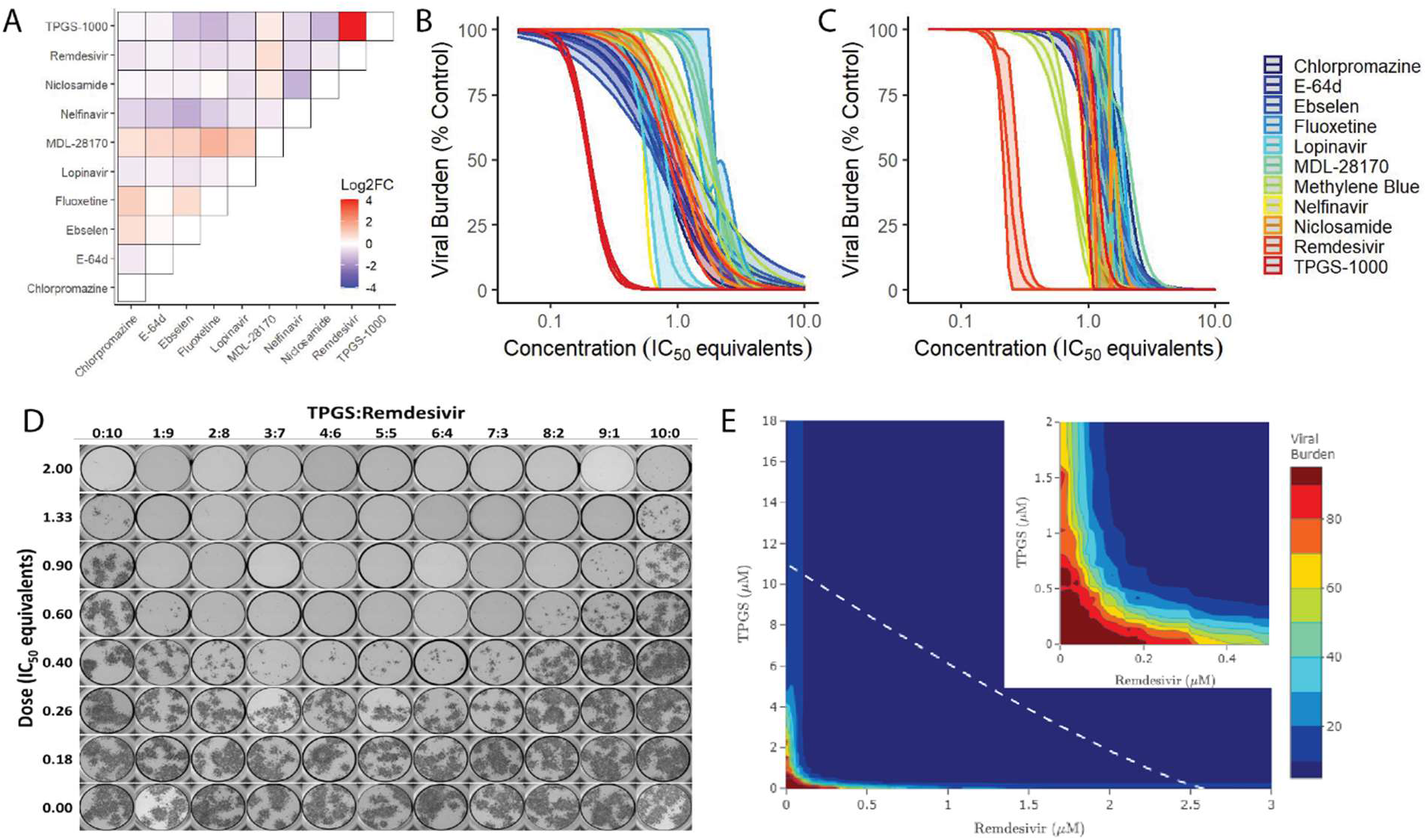
TPGS synergizes with remdesivir. A) Heatmap of the log2 fold change in potency for indicated combinations of the top results from screening. Values determined from three technical replicates. B,C) Curve-shift analysis for top drugs combined with B) Remdesivir or C) TPGS. Shaded regions surrounding each curve represent the 95% confidence interval for the dose-response model. D) Representative whole-well images captured for E) response-surface analysis of combinations of remdesivir and TPGS. The white, dashed line represents the predicted isobole for the IC_90_ while the inset magnifies the origin of the graph. Two technical replicates were used to determine the antiviral activity at each point assayed. All experiments were performed at least twice.

### α-Tocopherol drives TPGS activity

Given the pronounced synergy between remdesivir and TPGS, we next sought to understand the mechanism for the antiviral activity of TPGS, starting with its known, biologically-relevant activities: a non-competitive inhibitor of the drug exporter P-glycoprotein (PGP)^26, 27^, a surfactant, and a derivative of Vitamin E. Upon querying our list of screened drugs for substrates or inhibitors of PGP, we found no clear relationship between substrates (i.e. competitive inhibitors) of PGP and antiviral activity (Figure S5A). Additionally, we further screened well-characterized inhibitors and inducers of PGP for antiviral activity against SARS-CoV-2 (Figure S5B-G). While the majority showed no effect, amiodarone, another non-competitive inhibitor of PGP, exhibited antiviral activity against SARS-CoV-2 (Figure S5G). Amiodarone, however, showed no ability to synergize with remdesivir (Figure S4H), suggesting that the antiviral activities of TPGS and amiodarone are mediated by independent mechanisms and are therefore unlikely to be related to their commonality as PGP inhibitors. Additionally, TPGS is a surfactant. Other surfactant molecules with related chemical structures showed no antiviral activity (Figure S5I-L).

Finally, we tested the constituent components of TPGS: D-α-tocopherol, succinate, and polyethylene glycol with a molecular weight of 1000 Da (PEG-1000; Figure S6A). α-tocopherol alone is insoluble in an aqueous environment, so we instead tested α-tocopherol succinate (αTOS) and α-tocopherol phosphate (αTOP). While we observed no effect from αTOP, PEG-1000, or succinate alone (Figure S6B-D), we found that αTOS is capable of inhibiting SARS-CoV-2 replication in VeroE6 cells, albeit at higher concentrations than TPGS (IC_50_ = 21.7 µM; Figure S5E). These findings suggests that α-tocopherol is the active component of TPGS, and succinate and PEG-1000 contribute indirectly to its potency.

To determine whether the synergy with remdesivir and the increased potency of TPGS was specific to PEG-1000 or limited to the D-isomer of α-tocopherol, we tested a related compound: DL-α-tocopherol methoxypolyethylene glycol succinate (TPGS-750-M). TPGS-750-M is composed of an α-TOS molecule modified with a shorter PEG that terminates in a methoxy group, and the DL indicates that this compound consists of a racemic mixture of equal parts of the eight possible stereoisomers of α-tocopherol. Despite these differences, TPGS-750-M demonstrated comparable antiviral activity to TPGS (Figure S6F) as well as the ability to synergize with remdesivir (Figure S6G,H). Within logical constraints, these data suggest that medicinal chemistry targeting the size of the PEG or the isomer of the α-tocopherol may be modified to yield a compound with different physical and chemical properties that retains its antiviral activity.

### Tocopherols inhibit SARS-CoV-2 RdRp

We next sought to determine where in the process of viral propagation TPGS acts. We therefore infected VeroE6 cells with SARS-CoV-2 for 30 minutes and measured both intracellular and extracellular SARS-CoV-2 genome expression at 0, 3, and 9 hours post infection (hpi). We observed no decrease in viral attachment or uptake, but we detected a ∼4000-fold reduction in genomic material at 9 hpi in cells treated by either remdesivir or TPGS (Figure 3A). Notably, there was no detectable extracellular genome at any of these timepoints, indicating inhibition of viral transcriptional activity alone (data not shown).

**Figure 3.**
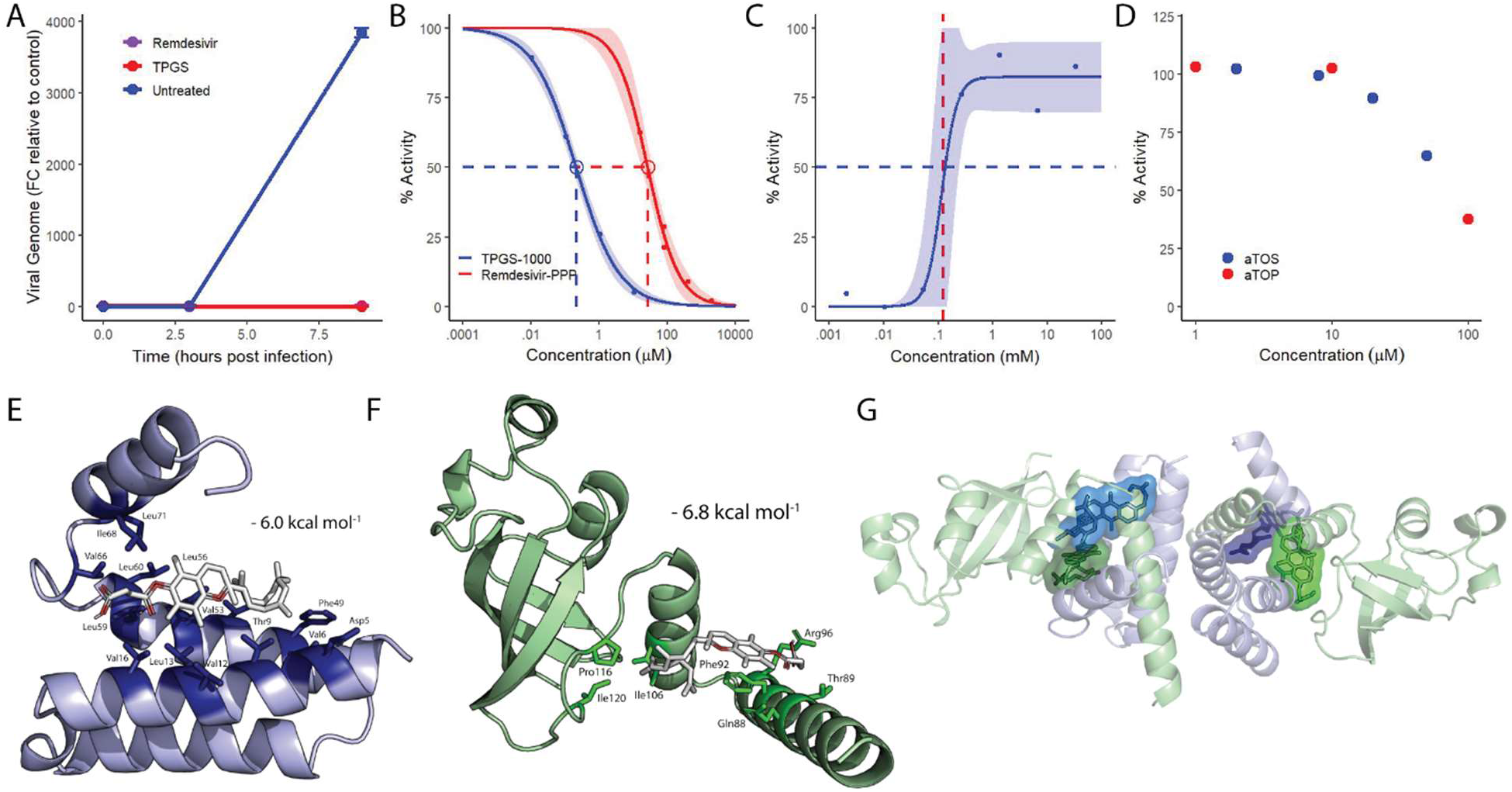
Water-soluble tocopherols inhibit the transcriptional activity of SARS-CoV-2 RNA-dependent RNA polymerase. A) Graph representing the fold change of SARS-CoV-2 genome in infected VeroE6 cells relative to initial uptake in the untreated control. Remdesivir and TPGS were given at a fully effective dose, 10 µM and 30 µM, respectively. Points and error bars represent the mean +/- SEM calculated from two technical replicates. B) Dose-response curves for the transcriptional activity of the SARS-CoV-2 replication complex treated with TPGS (blue) and remdesivir (red). The shaded region represents the 95% CI and points represent replicates from a single experiment. C) Dose-response curve for the transcriptional activity of the SARS-CoV-2 replication complex treated with high concentrations of TPGS, with 50% activity (blue, dashed line) and the critical micelle concentration of TPGS (red, dashed line) indicated. The upper limit of the model was not fixed. Points represent replicates from a single experiment. D) Dose-response plot for increasing concentrations of αTOS (blue) and αTOP (red). Points represent replicates from a single experiment. E) Docking model of the most favorable pose for αTOS (white) interacting with conserved, hydrophobic residues (dark blue, named) within NSP7 (light blue). F) Docking model of the most favorable pose for αTOS (white) interacting with conserved, hydrophobic residues (dark green, named) within NSP8 (light green). G) Representation of the heterotetrameric structure of NSP7 (light blue) and NSP8 (light green) with the most favorable poses of αTOS interacting individually with each NSP7 (dark blue) and NSP 8 (dark green) superimposed.

We expanded our analysis of the expression of SARS-CoV-2 RNA by sequencing mRNA from the samples in our time course experiment and analyzing the transcriptome of infected host cells. We observed the same profound reduction in transcripts mapped to each of the genes in the SARS-CoV-2 genome and detected no RNA corresponding to many of the viral genes in our treated samples (Figure S7A). The latter point, in particular, suggests that TPGS inhibits the transcription of viral genes. With regard to the transcription of host mRNA, principle component analysis revealed treated and untreated samples at 9 hpi have distinct transcriptomes (Figure S7B). Follow up gene ontology analysis of TPGS-treated and untreated cells at 9 hpi revealed that of the top 20 biological processes identified as enriched, ten were related to lipid metabolism (Figure S7C). The latter suggests that TPGS-treated cells respond to the tocopherol component of TPGS.

Considering the potent synergy between TPGS and remdesivir, a known inhibitor of the SARS-CoV-2 RNA-dependent RNA polymerase (RdRp)^28, 29^, as well as the results from our time course experiment, we hypothesized that TPGS inhibits the SARS-CoV-2 RdRp. To directly test our hypothesis, we measured the ability of TPGS to inhibit the transcriptional activity of purified SARS-CoV-2 RdRp composed of the catalytic subunit NSP12, and two accessory proteins, NSP7 and NSP8. Indeed, TPGS is a potent inhibitor of the transcriptional activity of the SARS-CoV-2 RdRp (IC_50_ = 212 nM; Figure 3B)—approximately 100-fold more potent than the active metabolite of remdesivir (RTP; IC_50_ = 26 µM; Figure 3B).

Interestingly, we also observed that TPGS does not inhibit RdRp at concentrations above its critical micelle concentration (CMC; Figure 3C)^30^. This led us to the expanded hypothesis that water-soluble derivatives of α-tocopherol inhibit the SARS-CoV-2 RdRp. We tested this hypothesis using αTOS and αTOP in our RdRp transcriptional assay. Consistent with our results in VeroE6 cells, αTOS was capable of inhibiting transcription; however, we also observed comparable inhibition of the SARS-CoV-2 RdRp by αTOP (Figure 3D). These findings suggests that while the efficacy of these compounds as antivirals may vary in the more complex cellular environment, water-soluble tocopherols inhibit the SARS-CoV-2 RdRp.

Finally, to further elucidate the interaction between water-soluble tocopherols and the SARS-CoV-2 RdRp, a series of computational docking studies were performed. Due to the challenge of modeling the flexibility of the PEG in the TPGS molecule as well as the ability of αTOS to inhibit the purified RdRp as well as viral replication in cells, we performed these studies for αTOS. Our docking studies across the individual components of the RdRp identified high affinity binding sites for αTOS within each NSP7, NSP8, and NSP12 (Figure 3E,F; Figure S8). Interestingly, while the top poses for αTOS were spread to a few locations across the surface of NSP12, our docking studies on the interaction of αTOS with NSP7 and NSP8 each converged to a single region (Figure S8A-C). In each case, the most energetically favorable binding poses localized with residues that participate in one of the two recently described hydrophobic interfaces that is required for the assembly of the functional RdRp (Figure 3G; Figure S8D)^6^. The residues within this interface are highly conserved across coronaviruses, and site directed mutagenesis of several residues within this interface reduced or ablated the transcriptional activity of the SARS-CoV-2 RdRp^6^. Taken together, these findings strongly support a mechanism by which TPGS prevents or destabilizes the assembly of the SARS-CoV-2 RdRp.

### TPGS promotes RNA capping by RdRp

Since TPGS is an inhibitor of the SARS-CoV-2 RdRp, we next hypothesized that the synergy of TPGS and remdesivir stems from independent mechanisms of inhibition of transcriptional activity. We combined TPGS and RTP at concentrations that independently led to intermediate inhibition of transcription in our RdRp assay and determined the combination index of these mixtures. Contrary to our hypothesis, we observed a purely additive combinatorial effect (Figure 4A). This suggests that the synergy between these compounds stems from a mechanism beyond the inhibition of transcription alone.

**Figure 4.**
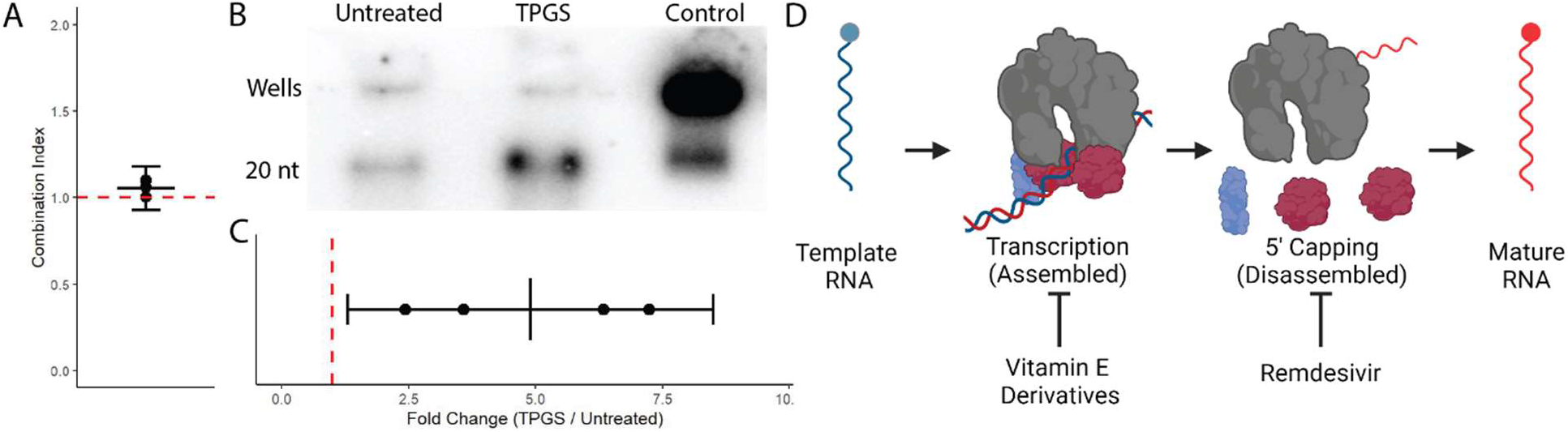
TPGS increases guanylyl transferase activity of SARS-CoV-2 RdRp. A) Plot of the combination index calculated for mixtures of TPGS and remdesivir at variable potency ratios (0.1 - 0.7 TPGS:remdesivir) and doses (0.4 – 1 IC_50_ equivalents). Each point represents the combination index determined in an independent experiment, the black line and error bars represent the mean +/- the 95% CI, and the red, dashed line represents a combination index of 1. B) Representative raw output from guanylyl transferase assay. C) Quantification of results from the guanylyl transferase assay pooling four replicates across three independent experiments. The black line and error bars represent the mean +/- the 95% CI and the red, and the red, dashed line represents a combination index of 1. D) Depiction of our proposed model for synergy between water-soluble derivatives of α-tocopherol and remdesivir.

Given these results and the recent description of remdesivir as an inhibitor of the guanylyl transferase (GTase) activity of NSP12^31^, we next decided to test GTase activity of NSP12 in the presence of TPGS. Compared to untreated, TPGS enhanced the GTase activity of the NSP12 in vitro (Figure 4B,C). This suggests that in the presence of TPGS, the RdRp function of NSP12 is inhibited, though the alternate GTase function of NSP12 is not. However, in the presence of both TPGS and remdesivir, both known activities of NSP12—essential for the production of mature viral RNA—are potently inhibited (Figure 4D). Blocking both distinct enzymatic functions could yield the observed synergy.

## Discussion

Here, we have identified water-soluble derivatives of tocopherol as inhibitors of the SARS-CoV-2 RdRp. Importantly, the most potent of these tested, TPGS, synergizes with remdesivir to yield a highly effective combination therapy. Docking studies suggest that tocopherol derivatives destabilize or inhibit the assembly of the SARS-CoV-2 RdRp by binding to hydrophobic residues at a highly conserved interface between NSP7 and NSP8 required for the assembly of the functional RdRp formed by NSP12, NSP7, and NSP8^6^.

While there are sites on the surface of NSP12 with marginally higher affinity for αTOS— most notably at the hydrophobic interface between NSP8 and NSP12—(Figure S8E), all top poses for the models of αTOS interacting with NSP7 and NSP8 localize to a single site within each protein, providing greater confidence in their relevance. This mechanism is further supported by the knowledge that these two sites together form a single interface required for the assembly of the functional SARS-CoV-2 RdRp^6^. Interestingly, this mechanism would explain our observation that TPGS treatment enhances the GTase activity of NSP12 in the presence of NSP7 and NSP8, as the disruption of the assembled RdRp complex in this assay would prevent NSP12 from participating in the assembled RdRp, thereby freeing all NSP12 to participate in 5’ capping.

We consider water-solubility a key factor in the potency of these compounds. Regardless of the mechanism, inhibition of the polymerase complex would require the tocopherol molecules to exist in a soluble form within the aqueous environment of the cell. Pegylation is a known method of solubilizing small molecules, and we attribute the considerable improvement in potency to PEG derivatization in TPGS when compared to αTOS. A corollary of this argument is that the antiviral activity of these compounds in a biological context relies on the stability of the solubilizing modification to the tocopherol which varies in biological systems^32^. Should the modification be hydrolyzed, the hydrophobic tocopherol will not remain in solution and will lose its antiviral effect. This may account for the variable efficacy of antiviral activity between αTOS and αTOP in viral replication assays (Figure S6D,E), despite comparable efficacy and potency in our transcriptional activity assay (Figure 3D). It is likely that both stability in an aqueous environment and hydrophobicity leading to greater cellular uptake both contribute to the antiviral activity of tocopherol derivatives tested here.

These considerations along with a vast literature highlight the potential of this group of molecules for optimization as clinical therapeutics. TPGS micelles have been studied extensively as a tool to enhance the solubility, oral-bioavailability, and the tissue penetration of numerous drugs^9, 33–36^. The critical micelle concentration of TPGS, as well as its capacity for PGP inhibition, can be shifted by modifying the length of the attached PEG molecule^37, 38^. The biological activity of Vitamin E is affected by its stereochemistry^39–41^, while its antiviral activity is not (Figure S6F). Finally, further solubilizing modifications should be explored to optimize solubility, potency, and stability, as well as the exploration of related tocopherols and isoprenoids. In combination or isolation, these characteristics can be modified to maximize antiviral efficacy while limiting potential host-directed effects of this direct-acting antiviral compound.

Though remdesivir is known to be a non-obligate terminator of transcription effective against SARS-CoV-2 and other viruses^28, 29, 42^, this activity alone does not account for the observed synergy—and therefore complete antiviral activity—observed when in combination with TPGS (Figure 4A). We note that remdesivir was recently described as an inhibitor of the GTase activity of NSP12^31^. TPGS does not inhibit this reaction; rather, the GTase activity of NSP12 is shown to be potentiated by TPGS (Figure 4B,C). We propose a model where TPGS and remdesivir synergize by inhibiting two sequential enzymatic reactions in the SARS-CoV-2 transcription cycle: polynucleotide synthesis and mRNA-capping, both of which are essential for cap-dependent translation in mammalian cells^43^. Presumably, both compounds also inhibit the two-step process of viral genomic replication, (+) and (-) strand synthesis of the coronavirus genome. Taken together, it is conceivable that this forms the conceptual framework for TPGS and remdesivir synergy. This model also provides an explanation for the consistent observation that the IC_50_ for the antiviral activity of remdesivir is orders of magnitude lower than the IC_50_ for the inhibition of the transcriptional or capping activity^2, 28, 44–46^. It follows, then, that the potency of remdesivir as an antiviral against SARS-CoV-2 may stem from its combined activity against both processes which can be enhanced by more potent transcriptional inhibition from TPGS.

In summary, we have identified a stable, safe, and orally-bioavailable compound, TPGS, that synergizes with the current standard of care, remdesivir, to inhibit SARS-CoV-2 replication. These findings also argue that inhibition of the GTase activity of NSP12 is an important part of the antiviral activity of remdesivir. Lastly, the antiviral activity of TPGS extends to a larger class of bioactive molecules with unique biophysical characteristics that retain their antiviral activity with chemical modification, opening the door to a new class of antiviral drugs.

## Methods

### Cell lines

Stocks of low-passage VeroE6 (ATCC # C1008) and Calu3 (ATCC # HTB-55) cells were stored in liquid nitrogen. As needed, stocks were revived and grown to confluency in a tissue-culture-treated flask at 37°C and 5% CO_2_ in complete minimal essential media (MEM) (Gibco # 11430-030) containing 10% fetal bovine serum (Denville Scientific Inc # C788U20); 2 mM glutamine (Gibco #25030-081); an antibiotic/antimycotic cocktail of peniciliin (100 units / mL), streptomycin (100 units / mL), and amphotericin B (250 ng / mL) (Gibco #15240-062); 25 mM HEPES (Gibco 15630-080); and 25 mM sodium bicarbonate (Gibco # 25080-094). At confluency, cells were detached using 0.25% trypsin-EDTA (Gibco # 25200-056), and seeded in a tissue-culture treated, 96-well microplate at a density of 4 x 10^4^ cells/well or dispensed to a flask (1:3) for maintenance. VeroE6 cells were seeded the day prior to experiments, and Calu3 cells were seeded four days prior to experiments.

### Virus

SARS-CoV-2 (Wa-1/USA) virus was obtained from BEI and propagated in VeroE6 cells. Virus was harvested from the cell culture supernatant by centrifugation at ∼ 1700 x g for 5 minutes to remove cell debris and subsequent collection of supernatant containing virus. The resulting preparation was aliquoted and stored at -80°C. All experiments in this manuscript were performed with third-passage or fourth-passage SARS-CoV-2 (10^7^ plaque forming units / mL). Sequence verification of SARS-CoV-2 WA-1/USA at each passage was performed with less than 10 synonymous invariants observed in each passage, and none in critical regions of the polymerase complex genetic loci.

### Drug Preparation and Storage

Information regarding the drugs used in screening experiments is available in Table S1. Where possible, drugs were prepared in 100% DMSO at a concentration of 20 mM, aliquoted to minimize freeze/thaw cycles, and stored at -20°C. If they were insoluble in DMSO, drugs were prepared in tissue culture grade water (Cytiva/Hyclone # SH30529.03). Drugs used in screening experiments were purchased from Millipore Sigma. PGP inhibitors and inducers in Figure S4 were purchased from Cayman Chemicals. Surfactants in Figure S4 were purchased from Millipore Sigma. The active metabolite of remdesivir, GS-443902, was purchased from Med Chem Express and stored at -80°C for a maximum of one month.

### Artificial-Intelligence-Driven Screen

The artificial intelligence tool mediKanren was used to efficiently identify FDA-approved, readily available drug compounds hypothesized to target human and/or viral proteins essential for SARS-CoV-2 infection for further screening in vitro (Figure S1). MediKanren is a biomedical reasoning system composed of a custom knowledge-graph database and a constraint-based logic reasoning engine, developed from the miniKanren family of relational-reasoning logic languages^47^. The reasoning engine finds biological relationships between biomedical concepts, such as molecules, drugs, proteins, pathways, genes, diseases, symptoms, and phenotypes and can identify candidates that are directly or indirectly connected. Its knowledge graph database incorporates over 180 distinct sources of biomedical knowledge, including the scientific literature (Pubmed), data on approved and unapproved drugs (via FDA and drugbank), and genes (NCBI, UniProt). This initial computational screen for human and/or viral proteins related to SARS-CoV infection produced approximately 1,300 compounds that were algorithmically prioritized and then manually investigated based on hypothesized therapeutic benefit. MediKanren was developed by the Hugh Kaul Precision Medicine Institute at UAB and source code for mediKanren software is available free of charge as open source on github (https://github.com/webyrd/mediKanren).

### Antiviral activity assay

One hour prior to infection, unless otherwise indicated, cells were washed once in serum-free (SF) MEM and incubated in SF-MEM containing the indicated treatment. Infections were performed for one hour in SF-MEM at 35°C at a multiplicity of infection (MOI) of 0.1 for SARS-CoV-2 in VeroE6 cells, an MOI of 1 for SARS-CoV-2 in Calu3 cells, and an MOI of 0.1 for OC43 in VeroE6 cells. Following the infection, an equal volume of media containing Avicel (FMC BioPolymer # Avicel CL-611) and fetal bovine serum was added to achieve a final concentration of 0.6% and 2%, respectively. Cells were then incubated at 35°C for 48 hours and fixed in 10% phosphate buffered formalin for a minimum of 1 hour and a maximum of 72 hours.

Following infection and fixation, cells were immunostained for SARS-CoV-2 spike protein. First, endogenous peroxidase activity in fixed monolayers was blocked by incubation in methanol containing 0.5% hydrogen peroxide for 30 minutes. Following three washes in deionized water, nonspecific protein binding was blocked by incubation in phosphate-buffered saline (PBS) containing 5% nonfat milk for 20 minutes. Cells were then directly incubated with a rabbit polyclonal antibody against SARS-CoV-2 spike/RBD protein (SinoBiological # 40592-T62), diluted 1:2500 in milk on a rocking platform for 1 hour at room temperature. Following this incubation, cells were washed five times with PBS and then incubated with horseradish-peroxidase-conjugated goat-anti-rabbit secondary antibody (Invitrogen # A16104) diluted 1:1000 in milk shaking for 1 hour. Cells were washed once in PBS containing 0.05% tween 20 and again five times with PBS. Finally, cells were incubated for 3 – 12 minutes with DAB reagent (Vector Laboratories), washed five times in water, and dried overnight while protected from light. For OC43, an identical procedure was followed, except that the primary antibody for viral detection was a monoclonal antibody against the coronavirus group antigen (Milipore Sigma #MAB9013) and the secondary was a horseradish-peroxidase-conjugated goat-anti-mouse antibody (enQuire BioReagents # Q2AB1).

Brightfield images of whole wells were acquired using either the Cytation 5 Cell Imaging Multi-Mode Reader (BioTek) or the Lionheart FX Automated Microscope (BioTek). Both instruments were operated with Gen5 software (BioTek), and images were acquired with the lowest power objective available—4x for the Cytation 5 and 1.25x for the Lionheart FX, respectively. Focusing was performed with the dual-peak (brightfield) method and stitching of images within a well was performed using the linear-blend method. Whole-well images were then processed in FIJI software^48^ where they underwent automated well-detection by thresholding and particle analysis followed by gaussian filtering, background flattening, and thresholding of the original image to determine the proportion of the well stained for viral protein.

The percentage of area determined to be positive for viral protein within each well was normalized to the mean value calculated across all control wells within a given experiment. This resulted in a standardized measurement of “viral burden”—a near-normally distributed variable with a high degree of concordance between replicates and experiments (Figure S2).

### Viability Assay

Uninfected VeroE6 cells were otherwise treated under the indicated condition as described in the antiviral activity assay. At 48 hours of treatment, cells were washed once in PBS and stained with Hoescht 33342 (ThermoFisher Scientific)—a membrane permeable nuclear stain—at 2 µg/mL and Sytox Green (ThermoFisher Scientific)—a membrane impermeable nuclear stain—at 1 µM for twenty minutes. Cells were washed three times with PBS and placed in PBS for imaging.

Automated imaging and analysis was performed using the Lionheart FX and Gen5 software. Hoescht 33342 and Sytox green staining was imaged with the 4x objective in two independent fields of view for each well, using the DAPI and GFP filter cubes, respectively. Acquired images were processed in Gen5 software by background flattening and blurring prior to nuclear segmentation based on Hoescht 33342 staining. Nuclear segmentation was performed by thresholding and watershed separation of touching objects. Nuclei below a conservative threshold for mean fluorescent intensity of Sytox green staining were considered live cells, and viability was calculated as a proportion of live cells to total cells within a well. 2000-4000 nuclei were analyzed per well.

### Statistical Analysis

All statistical analyses were performed with R statistical software (version 4.0.2)^49^ in the RStudio integrated development environment (version 1.3)^50^. Data were organized and graphed with the “tidyverse” collection of packages^51^, with the exception of response-surface analyses, which were performed with the “plotly” package^52^. Dose-response models predicting viral burden across a range of concentrations for a given intervention were calculated using the DRM function from the “DRC” package^53^. A four-parameter log-logistic model (equation 1) was used to fit viral burden to treatment dose:

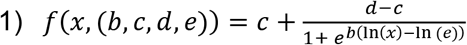

where *b* corresponds to the steepness of the curve, *c* is the lower asymptote of the curve, *d* is the upper asymptote of the curve, and *e* indicates the dose yielding a 50% effect. Because we operationally defined the untreated well as 100% viral burden and we modeled dose-response relationships only for drugs capable of suppressing viral burden to our limit of detection, *c* and *d* were fixed at 0 and 100, respectively, unless otherwise indicated. Gene Ontology analysis was performed with the “goseq” package as previously described^54^.

### Synergy analysis

The framework of Loewe additivity was applied for all assessments of synergy^24^. In brief, Loewe additivity relies on the dose equivalence principle—the dose of drug A that yields a given effect is equivalent to the dose of drug B that yields the same effect—and the principle of sham additivity—a drug added to itself is a purely additive effect^25^. Using the dose-response model for each drug (Figure 1B), we calculated the dose of the combined drugs in terms of IC50 equivalents as shown in equation 2^55^:

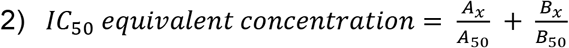

where *A*_*x*_ is the concentration of drug A that yields effect *x*, *A*_50_ is the IC_50_ for drug A, *B*_*x*_ is the concentration of drug B that yields effect *x*, and *B*_50_ is the IC_50_ for drug B.

For synergy screening, individual drugs were prepared at a concentration of 1 IC_50_ equivalent. Each compound was then mixed 1:1 with each other drug in a pairwise fashion, including itself (additive control), resulting in a mixture containing 0.5 IC_50_ equivalents of Drug A, 0.5 IC_50_ equivalents of Drug B, and a total dose of 1 IC_50_ equivalent. By definition, a purely additive combination of drugs given at this dose yields a 50% reduction in the viral burden for that well. Based on the principle of sham additivity, observed effects in additive control wells were used to determine the true dose administered for each drug in each experiment, and the expected additive effect for each mixture was adjusted accordingly. Given the potential for doses corresponding to the upper and lower portions of a dose response curve to lead to incorrect conclusions of antagonism and synergy, respectively^25, 56^, if the additive control for the given drug was outside of the linear range of the dose response curve (concentrations yielding approximately 20% -80% efficacy), all combinations of drugs containing that drug were excluded from analysis within that experiment.

For curve shift analysis, a similar approach to that described for synergy screening was used. In brief, drugs prepared at a concentration of 4 IC_50_ equivalents were combined 1:1 with another drug, resulting in an equipotent mixture of two drugs with a total dose of 4 IC50 equivalents. The resulting mixture was then serially diluted to yield a range of concentrations above and below the predicted IC_50_.

For isobologram and response surface analysis, the approach to curve shift analysis was expanded to encompass a range of potency ratios, also known as a fixed-ratio or “ray” design, as shown in equation 3^57^:

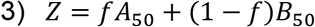

where *f* was increased from 0 to 1 by 0.1. This yields a series of mixtures, *Z*, ranging from 0% drug A and 100% drug B to 100% drug A and 0% drug B by 10% increments. Given that the potency ratio between two drugs may vary above and below their respective IC_50_ values, we represented the additive isobole for the IC_90_ as a curvilinear isobole, as shown in equation 4^58^:

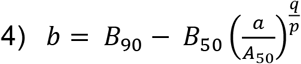

where *B*_90_ is the IC_90_ for drug B, *B*_50_ is the IC_50_ for drug B, a is a given concentration of drug A, *A*_50_ is the IC_50_ for drug A, q is the slope parameter calculated for the dose-response model of drug B, and p is the slope parameter calculated for the dose-response model of drug A.

### Viral uptake and replication assay

VeroE6 cells in a 6-well plate were infected with SARS-CoV-2 at an MOI of .1 for 30 minutes. At this point, considered 0 hpi, virus was removed, and cells were washed thoroughly with PBS three times and returned to complete MEM. At the indicated time points, cells were lysed and RNA was isolated using the Qiagen RNeasy minikit (Qiagen). RNA concentration was determined for each sample using a micro-drop spectrophotometer/fluorometer (DeNovix model DS-11 Fx +). The abundance of SARS-CoV-2 genetic material was subsequently determined by quantitative PCR using a Taqman probe and primers (Biosearch Technologies). The primers and probe sequences for the were as recommended by BEI resources for compatability with synthetic RNA standards (# NR-52358).

### mRNA sequencing and analysis

mRNA-sequencing was performed on the Illumina NextSeq500 as described by the manufacturer (Illumina Inc., San Diego, CA). Briefly, RNA quality was assessed using the Agilent 2100 Bioanalyzer. RNA with a RNA Integrity Number (RIN) of ≥7.0 was used for sequencing library preparation. RNA passing quality control was converted to a sequencing ready library using the NEBNext Ultra II Directional RNA library kit with polyA selection as per the manufacturer’s instructions (NEB, Ipswich, MA). The cDNA libraries were quantitated using qPCR in a Roche LightCycler 480 with the Kapa Biosystems kit for Illumina library quantitation (Kapa Biosystems, Woburn, MA) prior to cluster generation.

TrimGalore! (version 0.6.6) (https://www.bioinformatics.babraham.ac.uk/projects/trim_galore/) was used to trim the raw sequence FASTQ reads of primer adapter contamination (parameters used: --paired --trim-n --trim1 --nextseq 20). STAR (version 2.7.7a) was used to align the trimmed FASTQ reads to the combined Vero and SARS-CoV-2 Washington strain genomes (parameters used: --outReadsUnmapped Fastx --outSAMtype BAM SortedByCoordinate --outSAMattributes All)^59^. Following alignment, HTSeq-count (version 0.13.5) was used to count the number of reads mapping to each gene (parameters used: -m union -r pos -t exon -i gene_id -a 10 -s no -f bam)^60^.

### Transcriptional Activity of SARS-CoV-2 RNA-dependent RNA polymerase

The transcriptional activity of the SARS-CoV-2 RdRp was assessed using the SARS-CoV-2 RNA-dependent RNA polymerase kit plus (Profoldin #S2RPA100KE), according to the kit instructions for reactions in a 384-well plate. In brief, reactions were carried out at 35°C for 2 hours in a total of 25 µL in the presence of the indicated treatment condition. 65 µL of fluorescent dye were added and fluorescence was determined on a per-well basis in a Synergy H1 multi-mode plate reader (BioTek). 0% transcriptional activity was defined by the no-enzyme control and 100% activity was defined by an untreated control.

### Docking Studies

Docking experiments were performed using Autodock Vina^61^. Individual NSP7 and NSP8 structures were isolated from the structure of the SARS-CoV-2 NSP7-NSP8 complex (PDB ID: 7JLT)^6^ while SARS-CoV-2 NSP12 was isolated from the SARS-CoV-2 RNA-dependent RNA polymerase structure (PDB ID: 6M71)^62^. These structures and α-tocopherol succinate were modeled in Autodock Tools with α-tocopherol succinate possessing 16 rotatable bonds. The search space was defined as a 36.75 x 38.25 x 37.50 Å box for NSP7, a 48.75 x 52.50 x 41.25 Å box for NSP8, and an 82.50 x 89.25 x 88.12 Å box for NSP12 and encompassed the entire structure as to not bias the docking. No residues were designated as flexible in any of the receptors. The ten lowest energy solutions were then analyzed, and figures were generated in Pymol using a combination of the pose with the lowest energy as well as all poses found. Autodock Vina employs a strategy of initiating multiple runs starting from random ligand conformations. Free energy of binding calculations are performed using the intermolecular component of the lowest-scoring conformation as calculated via AutoDock Vina’s scoring function^61^.

### NSP12 Guanylyl Transferase Activity Assay

The GTase activity assay was performed as previously described^45^. To generate diphosphorylated RNA for the GTase activity assay, 5 μM of 20-base model RNA (5’-AAUCUAUAAUAGCAUUAUCC-3’; ThermoFisher Scientific) was treated with 300 nM of purified recombinant SARS-Cov2 NSP13 with an N-terminal His-tag (Cayman Chemicals #30589) in the presence of 50 mM Tris pH-7.4, 5 mM MgCl2, and 10 mM KCl for 15 mins at 30°C, followed by heat inactivation of NSP13 at 70°C for 5 mins.

The GTase reaction was carried out by incubating 500 nM of SARS-Cov-2 RNA-dependent RNA polymerase (complex of NSP12, NSP7 and NSP8; ProFoldin #RDRP-100S2) with 1μM diphosphorylated 20mer RNA in the presence of 0.05 μM of α-32P-GTP (PerkinElmer #BLU006H), 5 mM MgCl2, and 10 mM KCl for 240 mins at 30°C. A positive control GTase reaction using 0.5 U / μL of the vaccinia capping enzyme (New England Biolabs #M2080S) in place of the SARS-CoV-2 RdRp was performed under similar conditions. The reaction was stopped by addition of loading dye consisting of 80% formamide, 10 mM EDTA, 0.01% Bromophenol Blue, 0.02% xylene Cyanol and 15% glycerol. The reaction products were resolved on 15% denaturing PAGE gel with 7 M Urea (Bio-Rad #3450091), and was exposed to a phosphor screen (Amersham) overnight. The phosphor screen was scanned on the Typhoon FLA-7000IP (GE Healthcare). Densitometric analysis of images was performed using the gel analysis tool available in Fiji^48^.

## Acknowledgements

We thank the Southeastern Biosafety Laboratory (SEBLAB), an NIAID supported Regional Biocontainment Facility (NIAID UC6 AI058599) and leadership (Justin Roth, Chad Dunaway, Moustafa Awaden, and Simone Seay) for assistance in developing procedures and protocols in support of BSL3 studies of SARS-CoV-2. The authors would like to acknowledge Paul Wolkowitz, Joel Glasgow, Blake Frey, and Patrick Molina for their thoughtful feedback as well as for enabling the safe conduct of this research. Research reported in this publication was supported by the UAB High Resolution Imaging Facility, the UAB Heflin Genomics Core (P30CA013148), and the UAB SEBLAB Alabama Birmingham.

---

**Figure S1.**
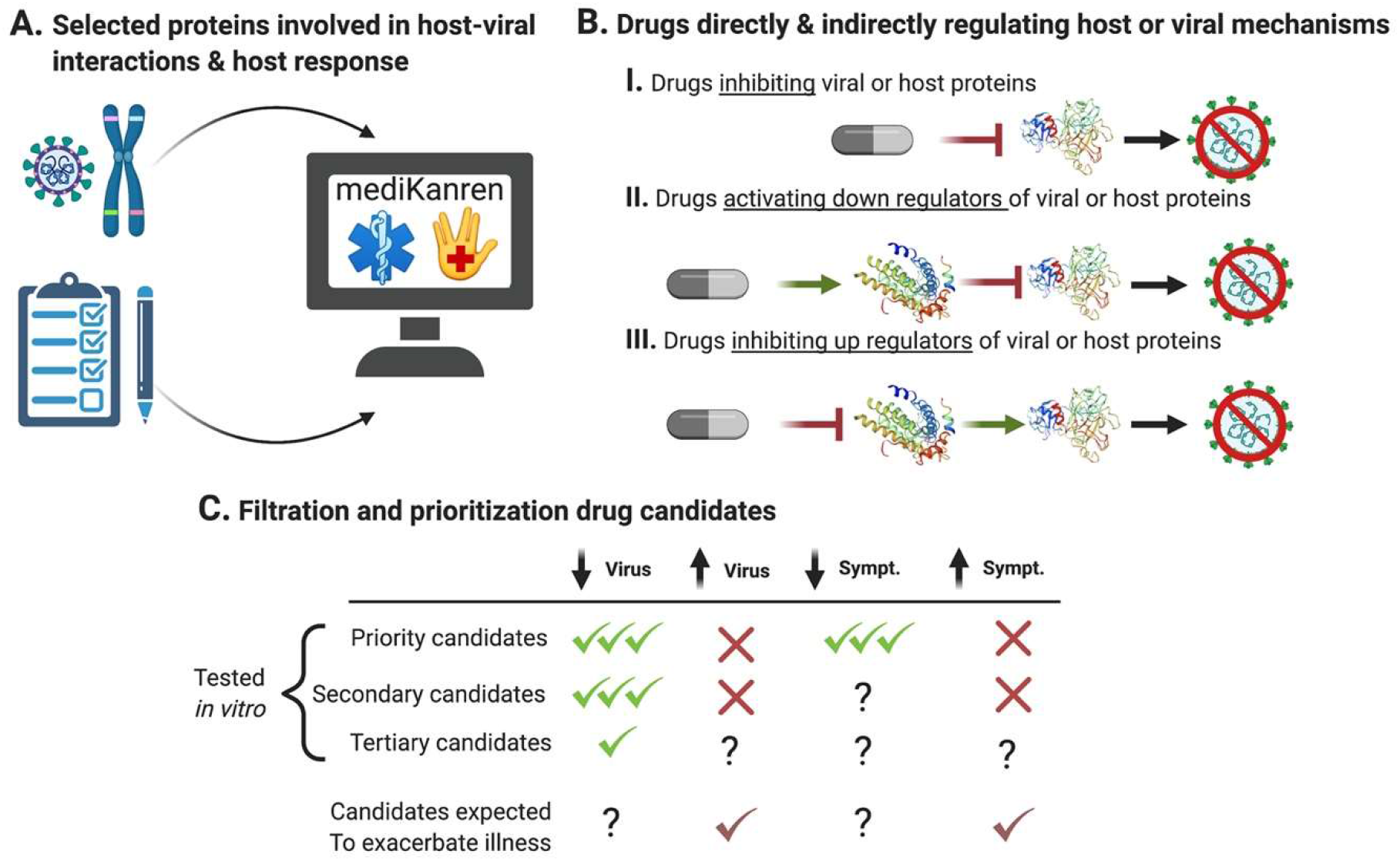
Depiction of expedited generation of candidate drugs based on host-pathogen interactions/pathways and prioritized based on therapeutic potential and risk. A) Host and viral proteins essential for viral replication and genes related to patients’ symptoms are input to mediKanren. B) Drugs with both direct and indirect impact on the protein’s activity are returned. C) Candidates are selected and given higher priority based on their ability to decrease mechanisms favorable to viral replication or those resulting in COVID-19 symptoms (impacting multiple mechanisms is shown as multiple check marks). A lower priority is given to candidates that would activate mechanisms favorable to viral replication or thought to increase COVID-19 symptoms. Question marks indicate ambiguous or lack of results. Candidates that are predicted to primarily increase viral replication or COVID-19 symptoms are flagged for review by the clinical trial committee. These may be included in observational trials to guide best practices for treating COVID-19 patients.

**Figure S2.**
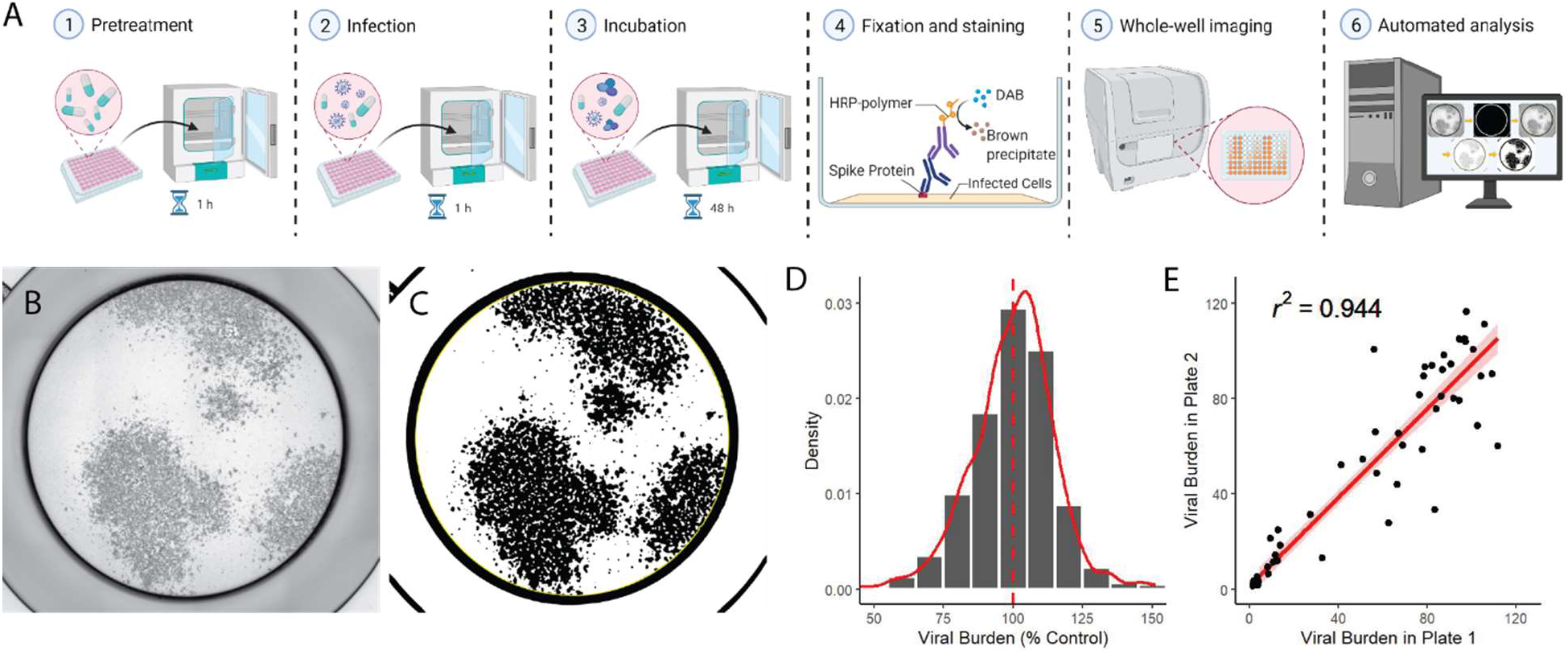
Focus-based antiviral activity assay is automated and reliable. A) An illustration depicting the major steps in the antiviral activity assay used for in vitro screening and follow up experiments. B) Raw and C) processed representative, whole well images from screening-based assay, including the automatically determined well area (yellow border). D) Histogram and density plot (red, solid line) from normalized, untreated control wells (n = 644) pooled across all experiments. The mean (red, dashed line) is mathematically centered on 100. E) Scatter plot of individual treatment conditions comparing the viral burden across replicates from two separate plates displaying the value for pearson’s correlation coefficient (r^2^), the linear regression line (red), and 95% CI (shaded region).

**Figure S3.**
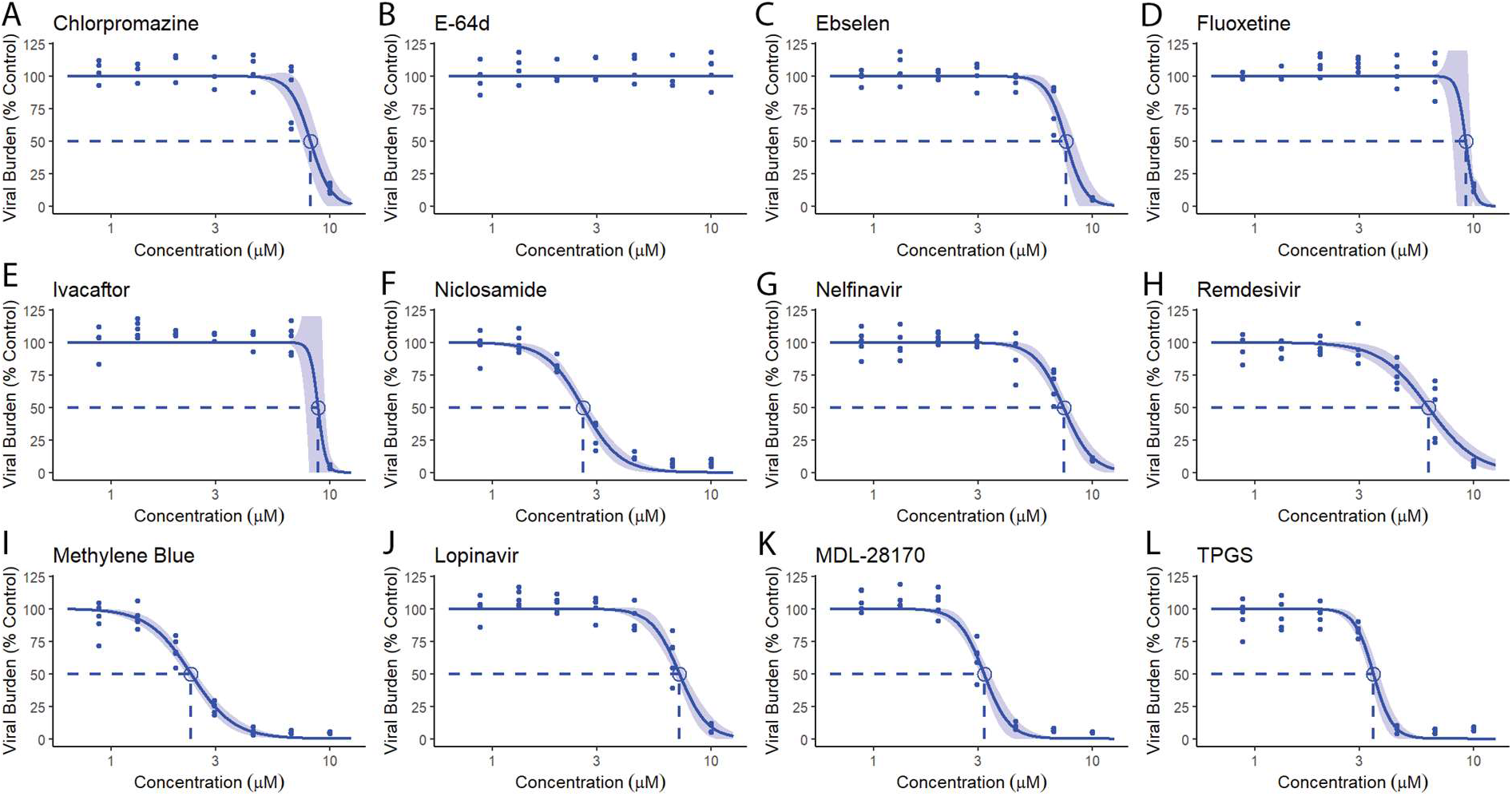
Top compounds inhibit β-coronavirus OC43. Dose response curves (solid line) and 95% CI (shaded region) for the antiviral activity of the top candidates against OC43. Points represent technical replicates pooled from two independent experiments, and the IC50 is indicated by the dashed lines and circle.

**Figure S4.**
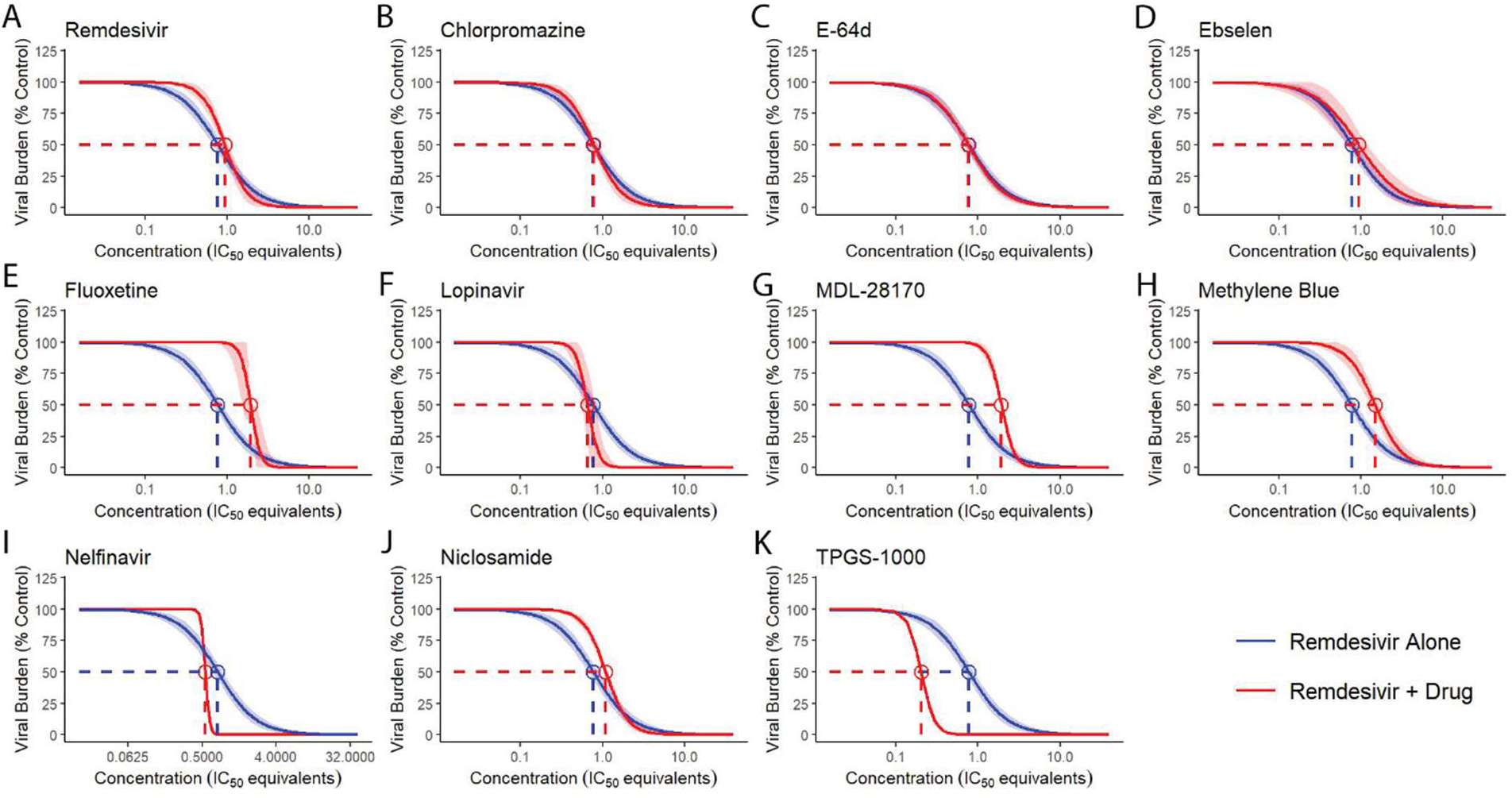
Most compounds combine with remdesivir in an additive fashion. Dose response curves (solid line) and 95% CI (shaded region) for remdesivir alone (blue) and remdesivir in an equipotent mixture with the indicated drug (red).

**Figure S5.**
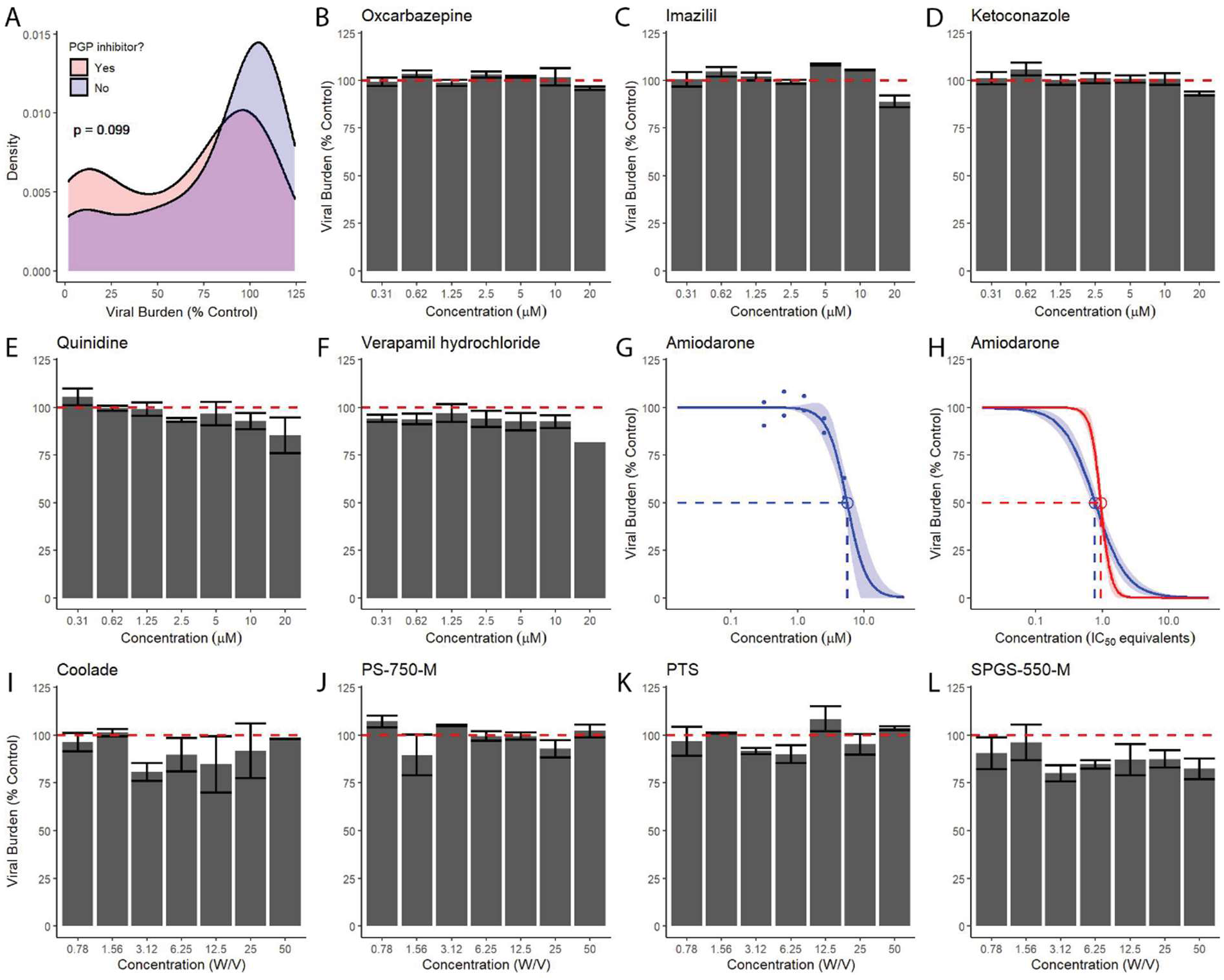
PGP-inhibition and surfactant activity do not account for the antiviral activity of TPGS. A) Density plots for screened drugs classified by DrugBank as inhibitors of PGP (red; n = 34) or not (blue; n = 63). Statistical significance determined by Kolmogorov-Smirnov test. B-F) Column graphs representing the mean +/- SEM for B) an inducer or C-F) inhibitors of PGP activity, determined by two technical replicates in a single experiment. G-H) Dose response curves (solid) and 95% CI (shaded) for G) Amiodarone, H) amiodarone plus remdesivir (red), or remdesivir alone (blue). Dots in G) represent individual replicates. I-L) Column graphs representing the mean +/- SEM determined by two technical replicates in a single experiment for surfactants with structural similarity to TPGS. A minimum of two independent experiments were performed for all panels.

**Figure S6.**
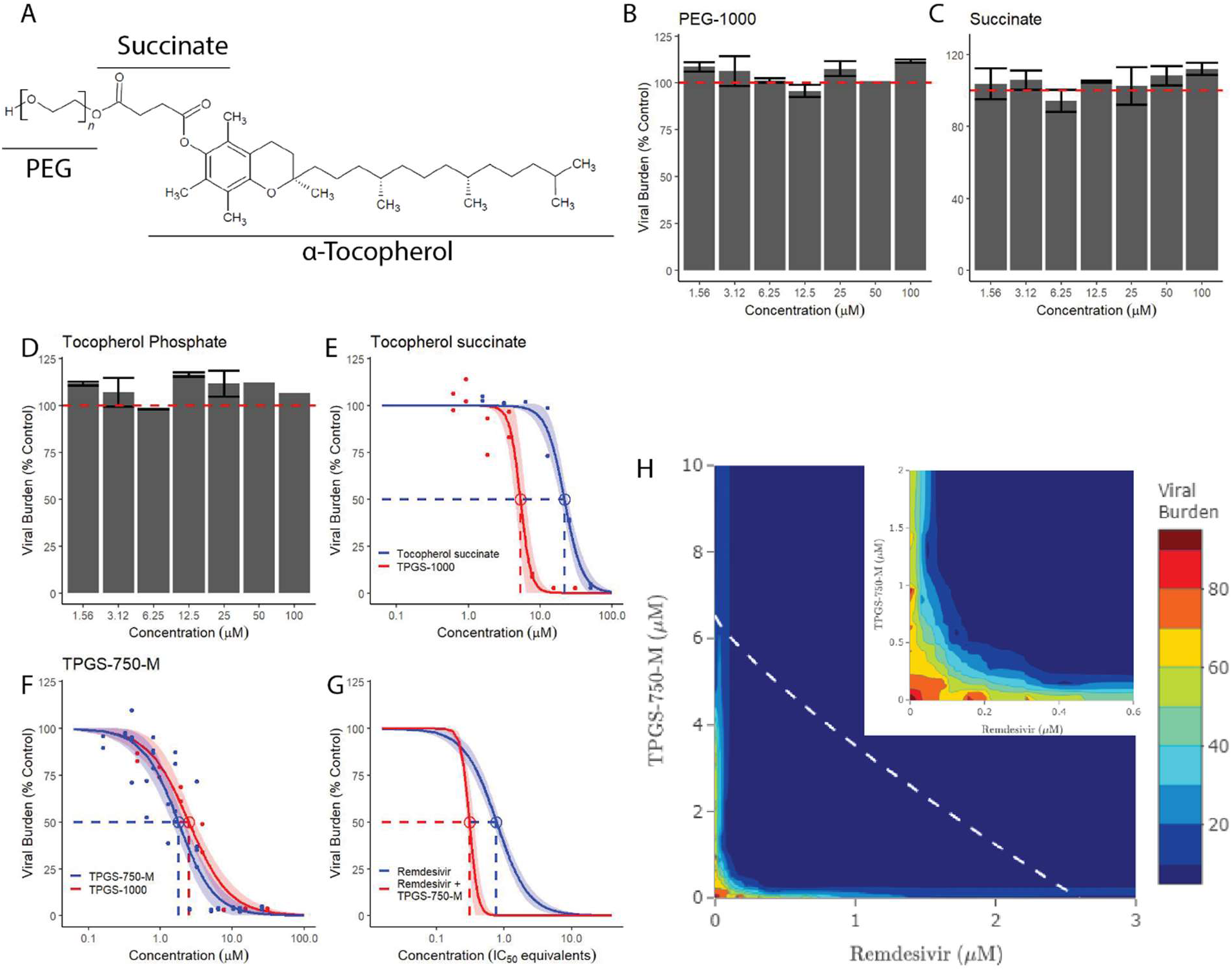
Water-soluble derivatives of Vitamin E inhibit SARS-CoV-2 replication. A) Molecular structure of a generic TPGS molecule. For αTOS, n = 0, for TPGS, n = ∼22. B-D) Column graphs representing the mean +/- SEM determined by two technical replicates in a single experiment for constituents of TPGS. E-G) Dose response curves (solid) and 95% CI (shaded) for E) αTOS (blue) and TPGS (red); F) TPGS-750-M (blue) and TPGS (red); and G) TPGS-750- M plus remdesivir (red) and remdesivir alone (blue). Dots in E,F) represent individual replicates. H) Response-surface analysis for TPGS-750-M in combination with remdesivir. The white, dashed line represents the predicted isobole for the IC_90_ while the inset magnifies the origin of the graph. Three technical replicates were used to determine the antiviral activity at each point assayed. A minimum of two independent experiments were performed for each panel.

**Figure S7.**
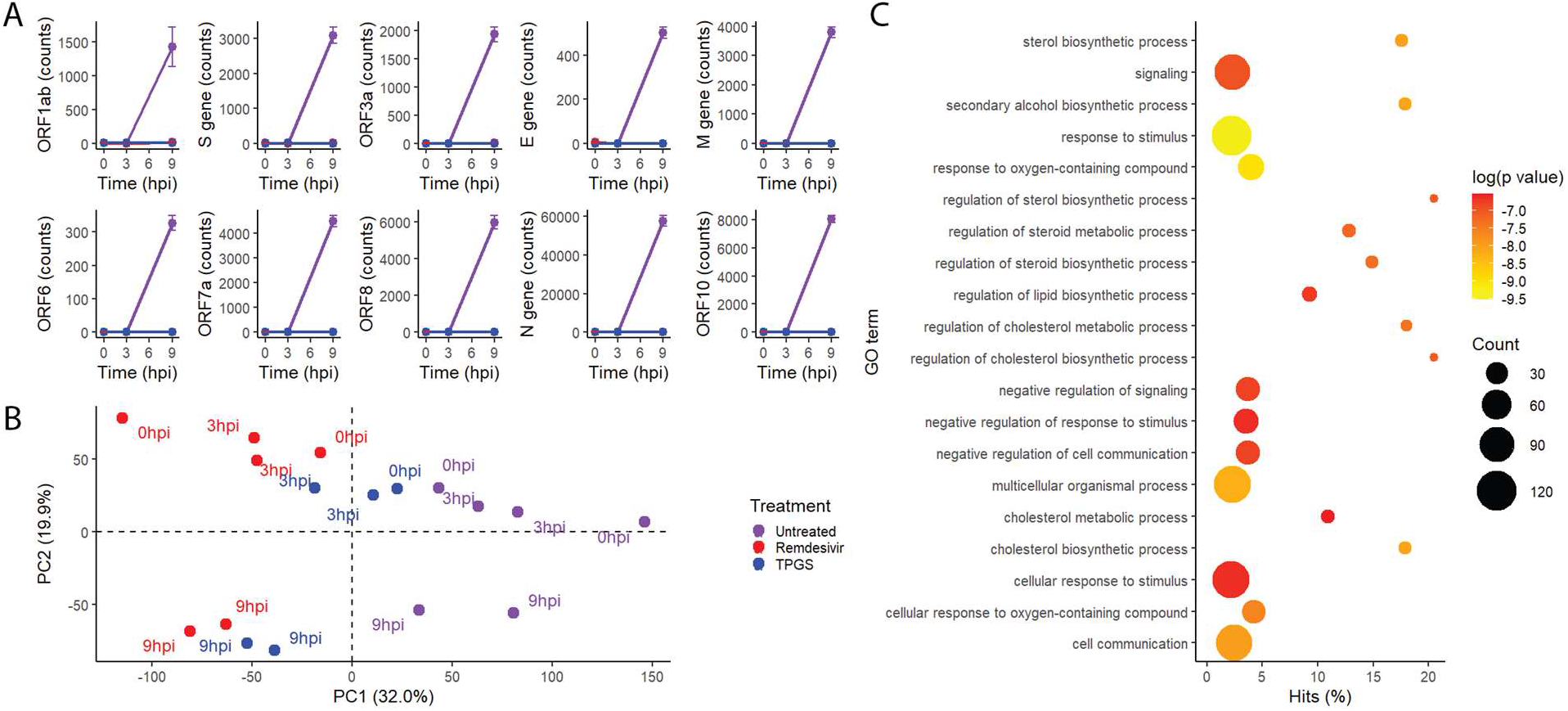
TPGS treatment reshapes the transcriptome of infected host cells. A) Graphs representing SARS-Cov-2 gene expression in infected VeroE6 cells. Remdesivir and TPGS were given at a fully effective dose, 10 µM and 30 µM, respectively. Points and error bars represent the mean +/- SEM calculated from two technical replicates. Group legend is in B). B) Principle component analysis of VeroE6 gene expression from the same samples depicted in (A). C) Analysis of the representation of genes with a Log_2_ fold increase greater than 1.5 in TPGS-treated samples at 9 hpi compared to untreated samples at 9 hpi among gene ontology categories.

**Figure S8.**
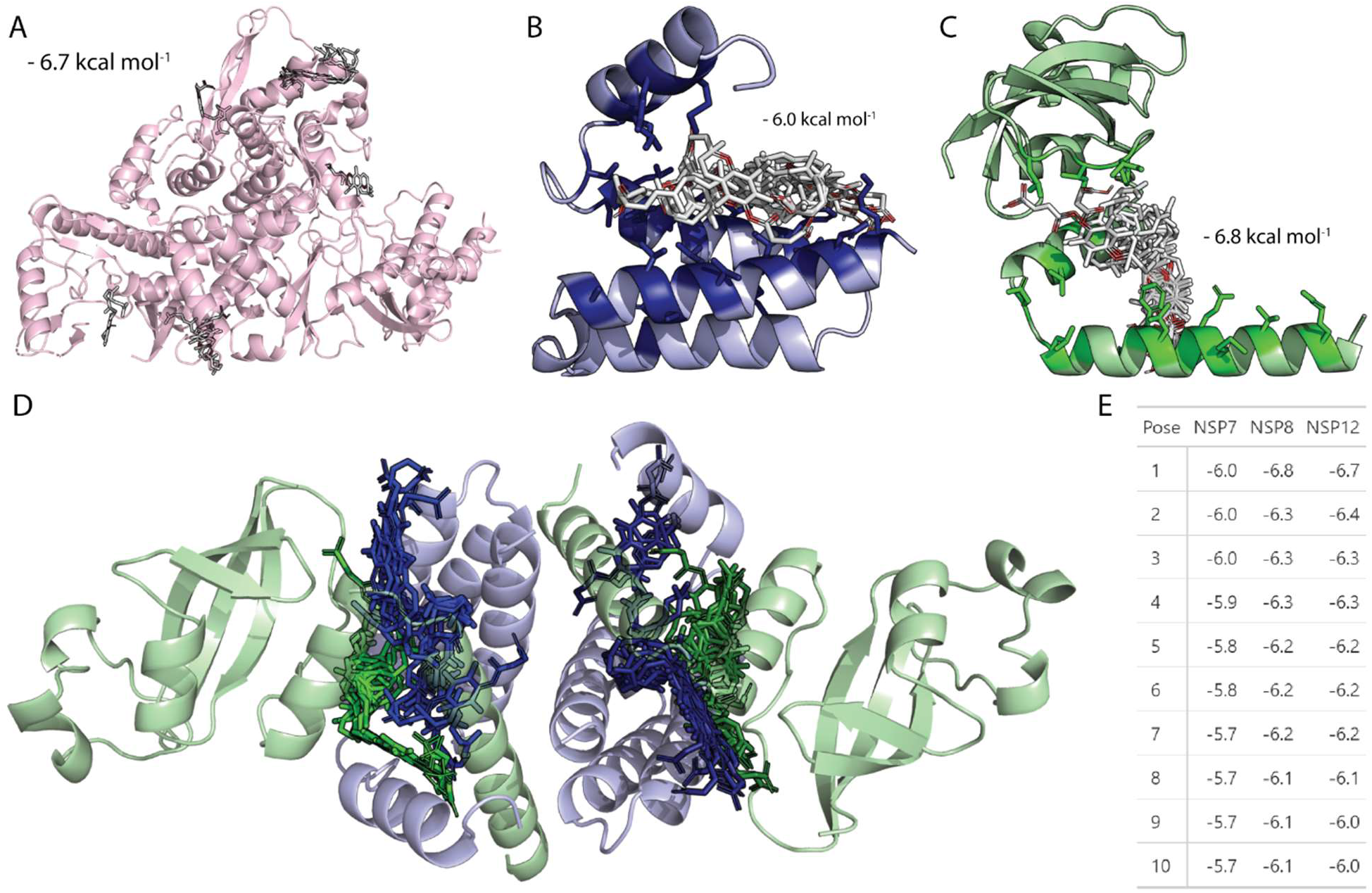
Binding of αTOS to conserved, hydrophobic interfaces in the SARS-CoV-2 RdRp is energetically favorable. A) Docking model of the ten most favorable poses for αTOS (white) interacting with NSP12 (light red). B) Docking model of the ten most favorable poses for αTOS (white) interacting with conserved, hydrophobic residues (dark blue) within NSP7 (light blue). C) Docking model of the ten most favorable poses for αTOS (white) interacting with conserved, hydrophobic residues (dark green) within NSP7 (light green). D) Representation of the heterotetrameric structure of NSP7 and NSP8 with the most favorable poses of αTOS (opaque) interacting individually with each NSP7 (transparent, blue) and NSP 8 (transparent, green) superimposed. E) Table of the top ten poses for αTOS interacting with NSP7, NSP8, and NSP12. Values represent the predicted binding affinity in kcal / mol.

## Notes

Funding: Supported by NIH grants R01HL149944 (KSH), R01AI134693 (CMP), U19 3U19AI142737 (TJG), T32HL13640 (HTP), Cystic Fibrosis Foundation grants HARROD19IO and HARROD20GO (KSH), UAB SOM COVID-19 Awards (CMP, MM, and KSH), Benjamin Monroe Carraway Endowment (KSH)

Competing Interests: The authors’ institution is pursuing a patent for the use of water-soluble tocopherol derivatives as antiviral compounds as described in this article.

### Competing Interest Statement

The authors institution is pursuing a patent for the use of water-soluble tocopherol derivatives as antiviral compounds as described in this article.

### Summary of Updates

Authors SA and AA were added to the author line.

